# Double-Negative B Cell Expansion Defines a Relapse-Prone Immunophenotype in Newly Diagnosed Giant Cell Arteritis

**DOI:** 10.64898/2026.09.25.754436

**Authors:** Beatriz Nieto-Carvalhal, Irene Monjo-Henry, Claudia Campos-Fabre, Sara García-Carazo, Cristina Bohórquez, Alejandro Villalba, Diana Peiteado, Eugenio de Miguel, María-Eugenia Miranda-Carús

## Abstract

**Background:** Giant cell arteritis (GCA), traditionally considered T cell-driven, exhibits vascular B/plasma-cell infiltrates and altered circulating B-cell homeostasis, suggesting a pathogenic role for B cells. Relapse is common in newly diagnosed GCA (nGCA), yet predictive biomarkers remain elusive.

**Objective:** To characterize circulating B-cell subpopulations in nGCA and assess their association with 12-month relapse.

**Methods:** Baseline multiparametric flow cytometry was performed on peripheral blood samples from consecutive patients with nGCA and sex/age-matched healthy controls (HC). Patients received standard treatment per 2018 EULAR recommendations and were followed longitudinally.

**Results:** 102 patients with nGCA and 102 matched HC were included. One-year outcome data were available for 55 patients, of whom 34 relapsed and 21 remained relapse-free. Baseline B-cell homeostasis differed by disease course. Non-relapsers showed reduced total CD19+ B-cell counts and contraction of most B-cell subsets, while CD27-IgD-double-negative (DN) B-cell counts remained comparable to HC. Conversely, relapsers maintained total B-cell numbers above HC despite reduced memory compartments, driven by expanded naïve and DN B cells, alongside preserved plasmablast and transitional B-cell counts. At 6 months, total B-cell counts increased only in non-relapsers. DN B cells showed the greatest divergence between groups and independently predicted relapse. DN B-cell counts also correlated with circulating T peripheral helper cell numbers, supporting extrafollicular immune pathway activation.

**Conclusion:** nGCA is characterized by outcome-dependent disturbances in B-cell homeostasis. Expansion of the DN B-cell compartment identifies a relapse-prone immunophenotype and represents a clinically relevant biomarker for early risk stratification.

**Key messages:**

- Outcome-dependent B-cell homeostasis distinguishes relapsing from non-relapsing newly diagnosed GCA.
- Baseline double-negative B-cell counts independently predict relapse after glucocorticoid monotherapy.
- Double-negative B-cell quantification may improve baseline risk stratification in GCA.

## INTRODUCTION

Giant cell arteritis (GCA), the most common systemic vasculitis in adults over 50 years of age [1–3], is marked by granulomatous inflammation of medium- and large-sized arteries [1–3], leading to tissue-destructive remodeling. Although its precise pathogenic mechanisms remain incompletely understood, immune dysregulation plays a central role, and GCA has traditionally been regarded as primarily T-cell driven [2]. Histologically, vascular lesions are characterized by infiltration of activated CD4⁺ T cells expressing high levels of PD-1 [2], together with macrophages and, less frequently, multinucleated giant cells [2]. Peripheral blood from patients with GCA shows increased proportions of circulating B-cell-helper T cell subsets [4–6], including PD-1^hi^ follicular helper (Tfh) and peripheral helper (Tph) CD4⁺ T cells [4–6]. More recent histopathological studies have demonstrated B cells and plasma cells within inflamed arterial tissue, both as diffuse infiltrates [7–9] and organized into tertiary lymphoid structures [4,7,8].

Beyond the vascular compartment, disturbances in circulating B-cell homeostasis have also been reported [9–14]. However, despite growing recognition of their potential relevance, the contribution of B cells to GCA pathogenesis remains incompletely defined. In particular, the extent to which changes in circulating B-cell populations are associated with disease activity and relapse risk remains unclear. In this context, we performed a prospective longitudinal study systematically characterizing circulating B-cell subpopulations in newly diagnosed GCA and evaluating their evolution under therapy as well as their potential association with subsequent relapse.

## PATIENTS AND METHODS

### Ethics Statement

The study was approved by the Hospital La Paz-IdiPAZ Ethics Committee, and all subjects provided written informed consent according to the Declaration of Helsinki.

### Patients

This is a prospective non-interventional study performed on consecutive patients referred to our ultrasound (US) GCA fast track clinic over a period of 24 months, in whom newly diagnosed GCA (nGCA) was clinically confirmed by the treating clinician after at least 6 months of follow-up (Table 1). Our fast-track clinic provides rapid clinical and US evaluation of patients referred from primary care and from La Paz University Hospital medical departments. In most cases Color Doppler US (CDUS) examination was performed within the first 24 h after referral, with a maximum of 72 h, and patients were naïve for glucocorticoids. Patients with autoimmune diseases other than GCA, cancer or concomitant infections were not included. Peripheral blood was drawn immediately upon initial diagnosis and after obtaining written informed consent. For each patient, an age and gender-matched healthy control (HC) was also studied. Subsequently, patients received standard glucocorticoid monotherapy per 2018 EULAR recommendations [15]. Close monitoring guided by our standardized GCA clinic protocol [6], enabled early detection of EULAR-defined relapses [15].

**Table 1.** Clinical Features of nGCA patients.

|  | **Clinical features (all)** | **Clinical features of patients with 1 year follow-up (n=55)** |  | **Relapse rate at 1yr attending to US/ clinical subgroups** |
| --- | --- | --- | --- | --- |
|  | **nGCA-all n=102** | **nGCA-nr n=34** | **nGCA-r n=21** | **Total Relapses n=21/55 (38%)** |
| Gender: F, n (%) | 58 F (57) | 23 F (67) | 14 F (66) |  |
| Age (yrs): median (IQR) | 81 (73-86) | 81 (77-85) | 82 (66-87) |  |
| Cranial, n (%) | 51 (50) | 19 (56) | 13 (62) | 13/32 (41) |
| Large Vessel, n (%) | 16 (16) | 7 (20) | 4 (20) | 4/11 (36) |
| Mixed, n (%) | 35 (34) | 8 (24) | 4 (19) | 4/12 (33) |
| Polymyalgia Rheumatica, n (%) | 36 (35) | 11 (32) | 9 (43) | 9/20 (45) |

### Color Doppler Ultrasound (CDUS)

Following established protocols [16–22], CDUS evaluated the superficial temporal (common trunk, frontal, parietal branches) and extracranial arteries (bilateral common carotid, axillary, subclavian). Based on these initial findings, patients were categorized into cranial (C), large-vessel (LV), or mixed (Mx) ultrasound phenotypes. Two expert rheumatologists performed all assessments (E.D.M. and I.M.H.).

### Temporal artery biopsy (TAB)

TAB was not performed routinely. Instead, it was reserved for clinically ambiguous cases, such as a positive CDUS with low pretest probability, or a halo sign restricted to a single arterial branch, according to 2018 EULAR recommendations [15].

### Immunophenotypic Analysis of Circulating B-Cell Subsets

The frequency and phenotype of B-cell subpopulations were assessed by flow cytometry of freshly isolated PBMCs. Events were acquired in a FacsCelesta flow cytometer with FacsDiva software (BD Biosciences) and analyzed with FlowJo 10.8.1 software. Fluorochrome-conjugated monoclonal antibodies were used to examine the expression of CD19 (BD, clone SJ25C1), CD20 (BD L27), CD27 (BD M-T271), CD24 (BD ML5), CD38 (BD HB7) and IgD (BD IA6-2). Absolute counts of CD19+ B cells and subpopulations were calculated using the Sysmex XN-9000 analyzer (Sysmex, Kobe, Japan) and dual-platform method. The detailed gating strategy for identifying circulating B-cell subpopulations is provided in Supplementary Figure S1. In addition, CD4+CXCR5-PD-1^hi^ peripheral helper cells (cTph) were identified as previously described [6, 23]; the detailed gating schema is reproduced in Supplementary Figure S2.

### Statistical Analysis

Comparisons between groups were performed using the Mann-Whitney or Kruskal-Wallis test followed by Dunn’s post hoc test for multiple comparisons. Paired samples were analyzed with the Wilcoxon test. Correlations were evaluated using Spearman’s rank correlation coefficients. Logistic regression was applied to verify the independent associations between immunological and clinical parameters. A receiver operating characteristic (ROC) analysis was performed to determine cutoff values with joint maximum sensitivity and specificity. All analyses were performed using Prism version 10.6.1 software for macOS (GraphPad, California, USA).

## RESULTS

### Patients

The baseline clinical characteristics of the 102 enrolled patients with nGCA are summarized in Table 1. At the time of analysis, first-year clinical outcome could be determined for 55 patients; among these, 21 (38.2%) experienced disease relapse (Table 1). According to the EULAR recommendations [15], these included 18 minor relapses and 3 major relapses. Relapse occurred after a median of 7 months (IQR 2-11) and at a median prednisone dose of 15 mg/day (IQR 7.5-22.5).

### Baseline total B-cell counts in nGCA

At baseline, patients with newly diagnosed Giant Cell Arteritis (nGCA, n = 102) exhibited significantly reduced total circulating B-cell counts compared with healthy controls (HC, n = 102) (Fig. 1A), showing only a weak correlation with baseline CRP and ESR levels (Fig. 1B). This reduction was consistently observed across all ultrasound-defined disease phenotypes (cranial, large-vessel, and mixed) and clinical subgroups, independent of concomitant polymyalgia rheumatica (PMR) (Fig. 1C). Notably, while the large-vessel vasculitis subgroup followed a similar downward trend, the difference did not reach statistical significance, likely reflecting the limited sample size of this cohort (Fig. 1C).

**Figure 1.**
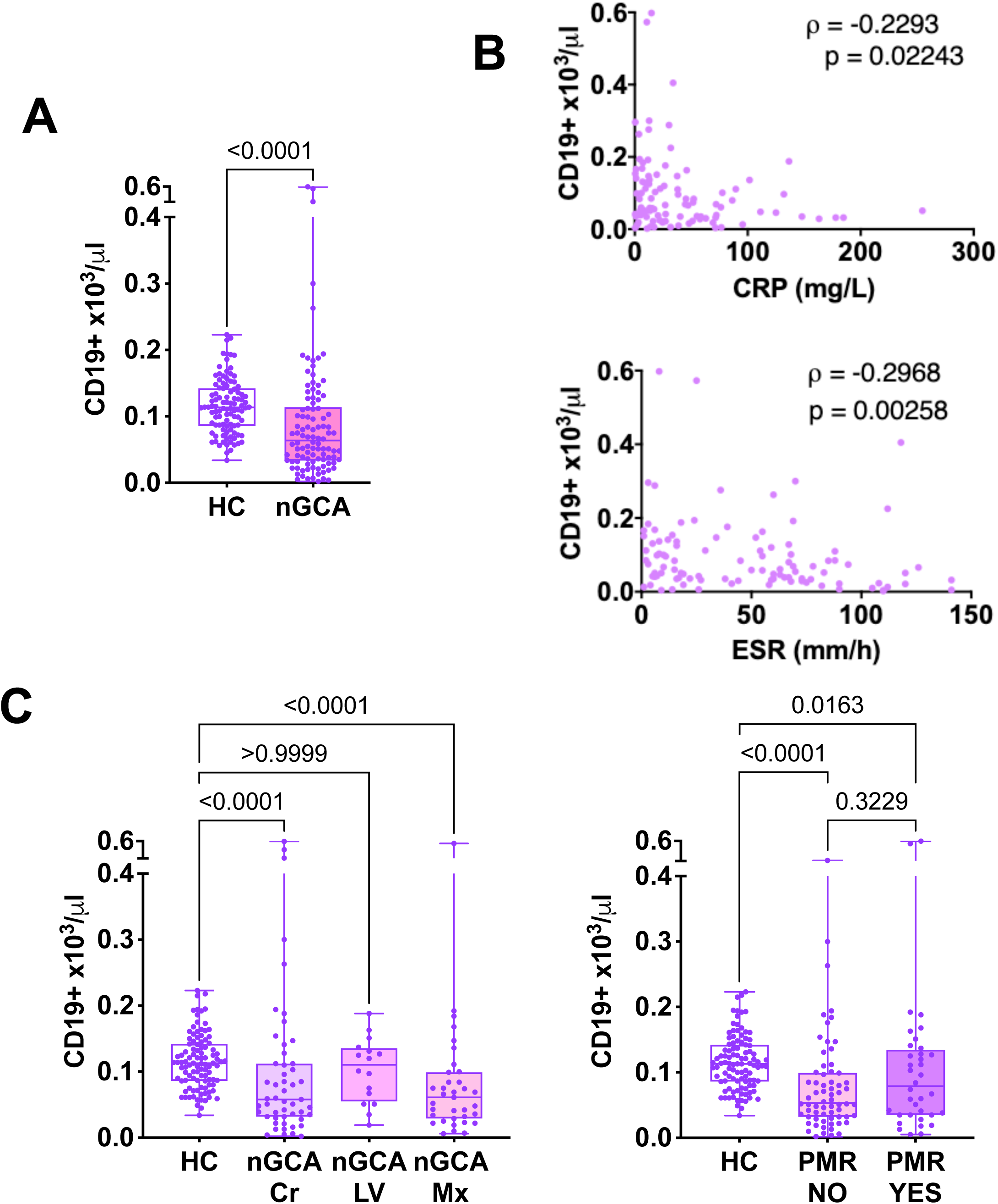
Baseline total B-cell counts in nGCA. (A) Baseline total B-cell counts in patients with nGCA (n = 102) and age- and sex-matched healthy controls (HC; n = 102). (B) Correlation of baseline B cell counts with C-reactive protein (CRP) and erythrocyte sedimentation rate (ESR). (C) Baseline total B-cell counts in nGCA patients stratified according to ultrasound-defined disease pattern [cranial (Cr), n = 51; large-vessel (LV), n = 16; mixed (Mx), n = 35] and the presence or absence of concomitant polymyalgia rheumatica (PMR) [PMR, n = 36; no PMR, n = 66]. P values for group comparisons were calculated using the Mann-Whitney U test. Correlations were assessed using Spearman’s rank correlation coefficient (ρ). **Alt text**. Patients with newly diagnosed giant cell arteritis exhibit reduced circulating B-cell counts compared with healthy controls, irrespective of ultrasound-defined disease pattern or concomitant polymyalgia rheumatica.

### Baseline B-cell subpopulation profile in nGCA

In addition to lower absolute counts, nGCA patients demonstrated a significantly altered B-cell subset distribution compared with HC (Fig. 2-A,B). This shift was characterized by decreased proportions of both unswitched (CD19+CD27+IgD+) and switched memory (CD19+CD27+IgD-) B cells (Fg. 2-A,B). Conversely, proportions of naïve (CD19+CD27-IgD+), double-negative (DN) B cells (CD19+CD27-IgD-) and plasmablasts (CD19+CD27+CD20-CD38^hi^) were significantly increased (Fig. 2-A,B). At the same time, the proportion of circulating transitional B cells (CD19+CD24^hi^CD38^hi^) was not different from that observed in HC (Fig. 2-A,B).

**Figure 2.**
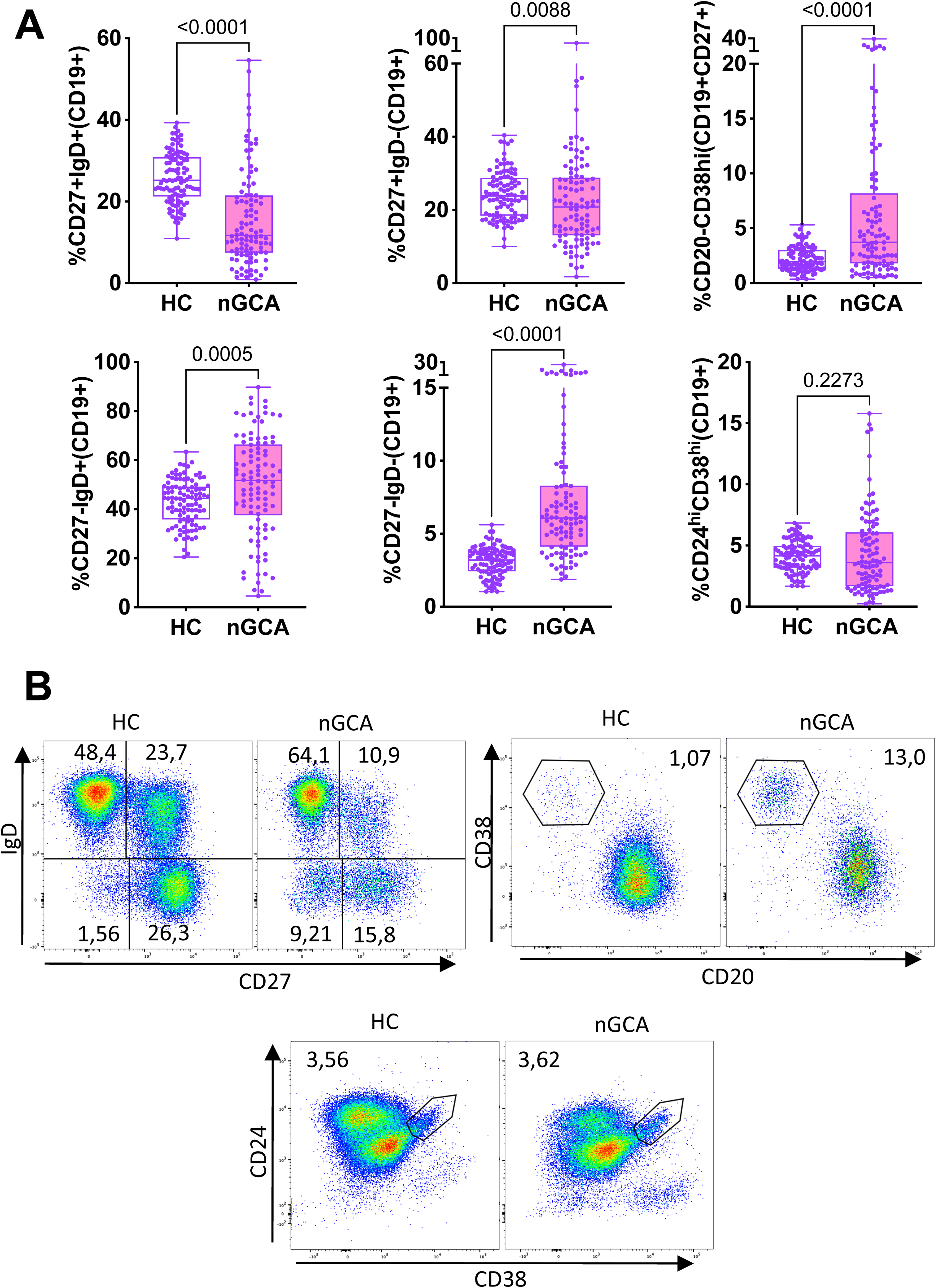
Baseline subpopulation profile in nGCA. (A). Frequencies of B-cell subsets are shown from left to right and top to bottom: unswitched memory B cells (CD19+CD27+IgD+); switched memory B cells (CD19+CD27+IgD-); plasmablasts (CD19+CD27+CD20-CD38^hi^); naïve B cells (CD19+CD27-IgD+); double-negative (DN) B cells (CD19+CD27-IgD-); and transitional B cells (CD19+CD24^hi^CD38^hi^). P values were calculated using the Mann-Whitney U test (B). Representative flow cytometry dot plots of CD27 and IgD expression in CD19+-gated cells (top left); CD20 and CD38 expression in CD19+CD27+-gated cells (top right) and CD24 and CD38 expression in CD19+-gated cells (bottom). **Alt text.** Patients with newly diagnosed giant cell arteritis display an altered circulating B-cell subset profile compared with healthy controls, characterized by reduced frequencies of memory B-cell subsets and expansion of naïve and double-negative B cells. Representative flow cytometry dot plots illustrate the gating strategy used to identify the analyzed B-cell populations.

### Longitudinal tracking of total B-cell counts and subpopulation profiles in nGCA

To assess the stability of the B-cell compartment, longitudinal changes in total B-cell counts were evaluated among 38 nGCA patients who provided a second blood sample 6 months after treatment initiation. At this time point, all patients had inactive disease while receiving prednisone (15-20 mg/day). Total B-cell counts had increased significantly from baseline across the entire cohort (Fig. 3A); however, stratification by 1-year relapse status revealed divergent patterns (Fig. 3A). This longitudinal increase was confined to patients who remained relapse-free (n = 23) (Fig. 3A), whereas those who relapsed (n = 15) exhibited no significant change in total B-cell counts (Fig. 3A). At the same time, ESR and CRP had normalized in both relapsers and non-relapsers (Fig. 3B).

**Figure 3.**
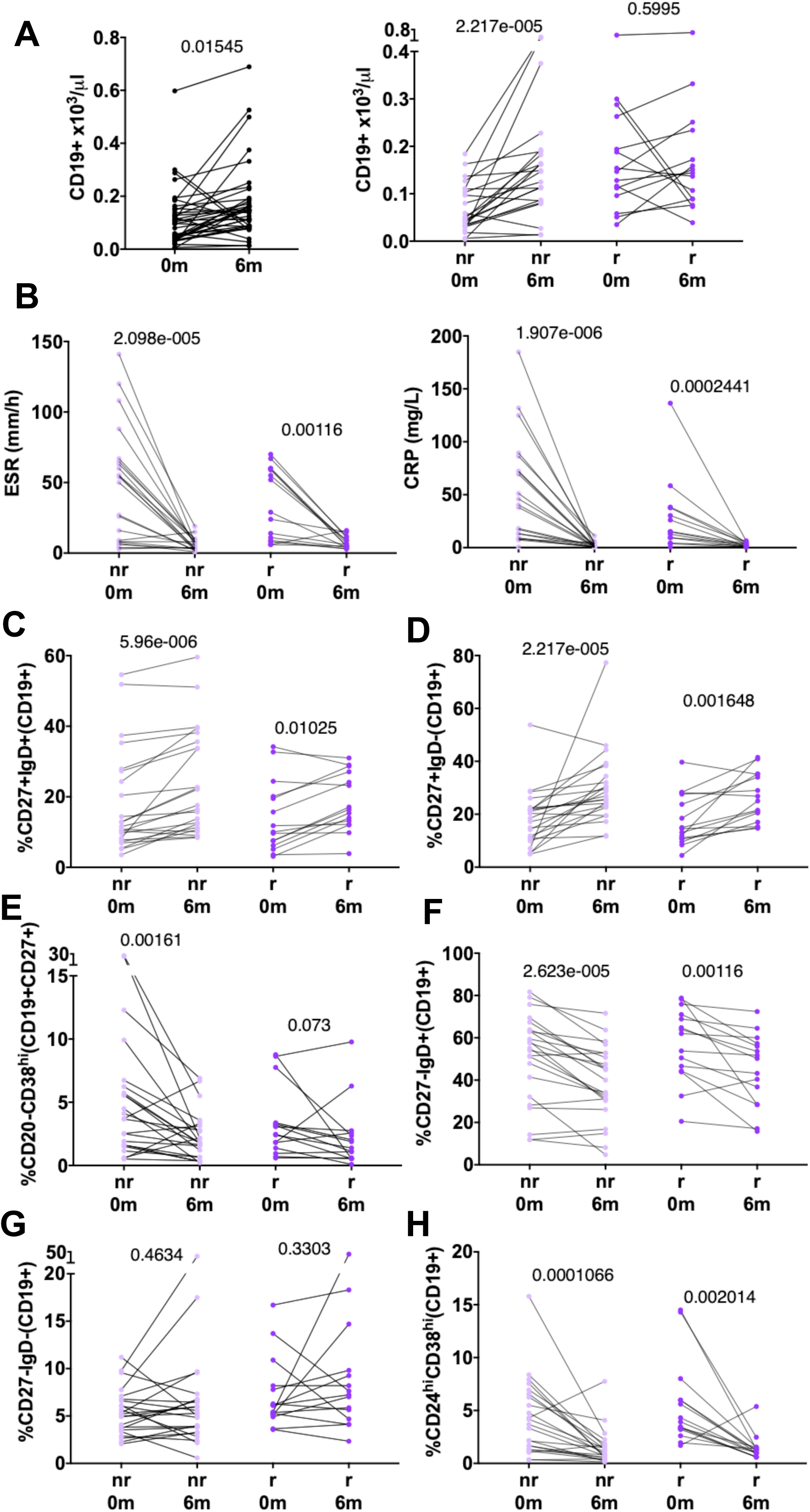
Longitudinal tracking of circulating B-cell populations and inflammatory markers during the first 6 months of glucocorticoid treatment. Paired longitudinal analyses comparing baseline (0 months) and 6-month (6 m) measurements in the subset of patients with available 6-month follow-up blood samples (n = 38). **(A)** Absolute CD19+ B-cell counts in this longitudinal cohort (n = 38, left) and stratified by their subsequent 1-year clinical outcome into non-relapsers (nr; n = 23) and relapsers (r; n = 15, right). **(B-H)**. Paired analyses stratified by clinical outcome: **(B)** Erythrocyte sedimentation rate (ESR) and C-reactive protein (CRP). **(C)** Unswitched memory B cells (CD19+CD27+IgD+); **(D)** Switched memory B cells (CD19+CD27+IgD-); **(E)** Plasmablasts (CD19+CD27+CD20-CD38^hi^); **(F)** Naïve B cells (CD19+CD27-IgD+); **(G)** Double-negative (DN) B cells (CD19+CD27-IgD-); and **(H)** Transitional B cells (CD19+CD24^hi^CD38^hi^). P values were calculated using the Wilcoxon signed-rank test. **Alt text.** Longitudinal tracking of B-cell populations and inflammatory markers. Paired baseline and 6-month analyses of total and stratified cohorts, showing B-cell subsets, ESR, and CRP by 1-year relapse status.

In parallel, analysis of B-cell subpopulation profiles revealed that both relapsers and non-relapsers demonstrated a significant increase in unswitched and switched memory B cell frequencies by month 6 (Fig.3-C,D). Concurrently, the frequencies of naïve B cells, transitional B cells, and plasmablasts decreased significantly in both clinical groups (Fig. 3-E,F,H). Conversely, despite these dynamic changes observed across the other B-cell subsets, the frequency of DN B cells did not change significantly in either group (Fig. 3G).

### Baseline B-cell counts according to clinical outcome

Following these longitudinal observations, we next investigated whether baseline differences in the B-cell compartment associate with relapse. Among the 55 patients with available 1-year outcome data, stratification by relapse status revealed a striking contrast (Fig. 4A). Compared with HC, non-relapsers (n = 34) exhibited significantly lower baseline B-cell counts (Fig. 4A), whereas relapsers (n = 21) displayed elevated levels (Fig. 4A), resulting in a marked separation between the two patient groups. (Fig. 4A). Notably, baseline ESR and CRP values were not significantly different between patients who subsequently relapsed and those who did not (Fig. 4B).

**Figure 4.**
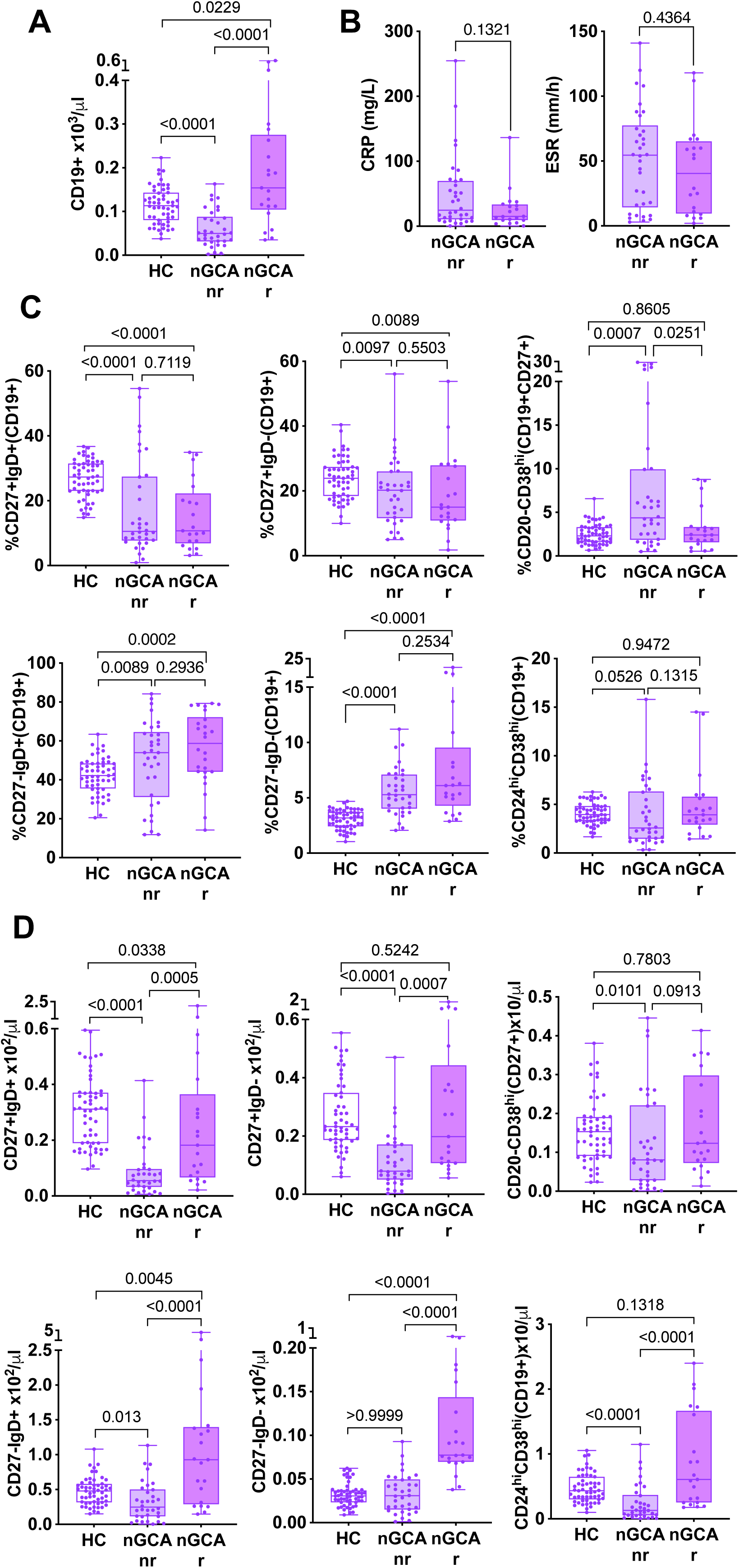
Baseline total B-cell counts and subset distribution in nGCA stratified by relapse within 12 months after diagnosis. Baseline measurements evaluating all patients with available 1-year clinical outcome data (n = 55) and healthy controls (HC; n = 55). (A) Baseline total B-cell counts in non-relapsers (nr; n = 34), relapsers (r; n = 21), and healthy controls (HC; n = 55). (B) Baseline C-reactive protein (CRP) and erythrocyte sedimentation rate (ESR) in non-relapsers and relapsers. (C) Baseline frequencies and (D) absolute counts of B-cell subsets in non-relapsers, relapsers, and healthy controls. Subsets are shown from left to right and top to bottom: unswitched memory B cells (CD19+CD27+IgD+); switched memory B cells (CD19+CD27+IgD-); plasmablasts (CD19+CD27+CD20-CD38^hi^); naïve B cells (CD19+CD27-IgD+); double-negative (DN) B cells (CD19+CD27-IgD-); and transitional B cells (CD19+CD24^hi^CD38^hi^). P values were calculated using the Kruskal-Wallis test with Dunn’s multiple-comparisons correction. **Alt text.** Baseline B-cell profiles differ according to 12-month clinical outcome in newly diagnosed giant cell arteritis. Patients who subsequently relapse exhibit higher total B-cell counts and expansion of naïve and double-negative B cells, whereas non-relapsers show reduced total B-cell counts and contraction of most B-cell subsets compared with healthy controls.

### Baseline B-cell subpopulation profile according to clinical outcome

Stratified analyses according to clinical outcome demonstrated altered baseline B-cell subset distributions in both patient groups relative to HC (Fig. 4-C,D). Compared with HC, both relapsers and non-relapsers exhibited significantly reduced proportions of unswitched and switched memory B cells, together with increased frequencies of naïve and DN B cells (Fig. 4C). No statistically significant differences were observed between the two clinical groups across most B-cell subsets (Fig. 4C). By contrast, a different pattern emerged upon analysis of absolute counts (Fig. 4D). Among non-relapsers, consistent with the reduction in total B-cell counts, the absolute numbers of most subpopulations were also reduced. DN B cells represented a notable exception, as their absolute counts were comparable to HC (Fig. 4D). In relapsers, despite the expansion of the total B-cell compartment, absolute unswitched memory B-cell counts remained significantly lower than in HC (Fig. 4D), whereas switched memory, transitional and plasmablast counts did not significantly differ from HC (Fig. 4D). In contrast, naïve and DN B-cell counts were significantly increased relative to HC (Fig. 4D). Consequently, relapsers exhibited significantly higher absolute counts of multiple B-cell subsets than non-relapsers (Fig. 4D), including non-switched memory, switched memory, naïve, DN and transitional B cells (Fig. 4D). Notably, the clearest distinction between the two groups was observed within the DN B-cell compartment (Fig. 4D), prompting us to evaluate its potential utility as a predictive biomarker.

### Association of baseline DN B-cell counts with relapse risk

Higher baseline DN B-cell counts were significantly associated with subsequent relapse [odds ratio (OR), 28; 95% confidence interval (CI), 12-51]. In multivariable logistic regression analysis, this association remained significant after adjustment for ESR, CRP, age, sex, ultrasound-defined subgroup, and the presence of PMR. Receiver operating characteristic (ROC) analysis demonstrated excellent discrimination for relapse (AUC, 0.93; 95% CI, 0.86-0.99) (Fig. 5A). The optimal cut-off value, determined using Youden’s index, was 0.06 DN B cells/µL. Values above this threshold indicated an increased risk of relapse, yielding a sensitivity of 86% and a specificity of 94%. Corresponding positive and negative likelihood ratios were 14.6 and 0.16, respectively, while the positive and negative predictive values were 90% and 91%. These findings support the potential utility of baseline DN B-cell counts as a predictive biomarker of relapse.

**Figure 5.**
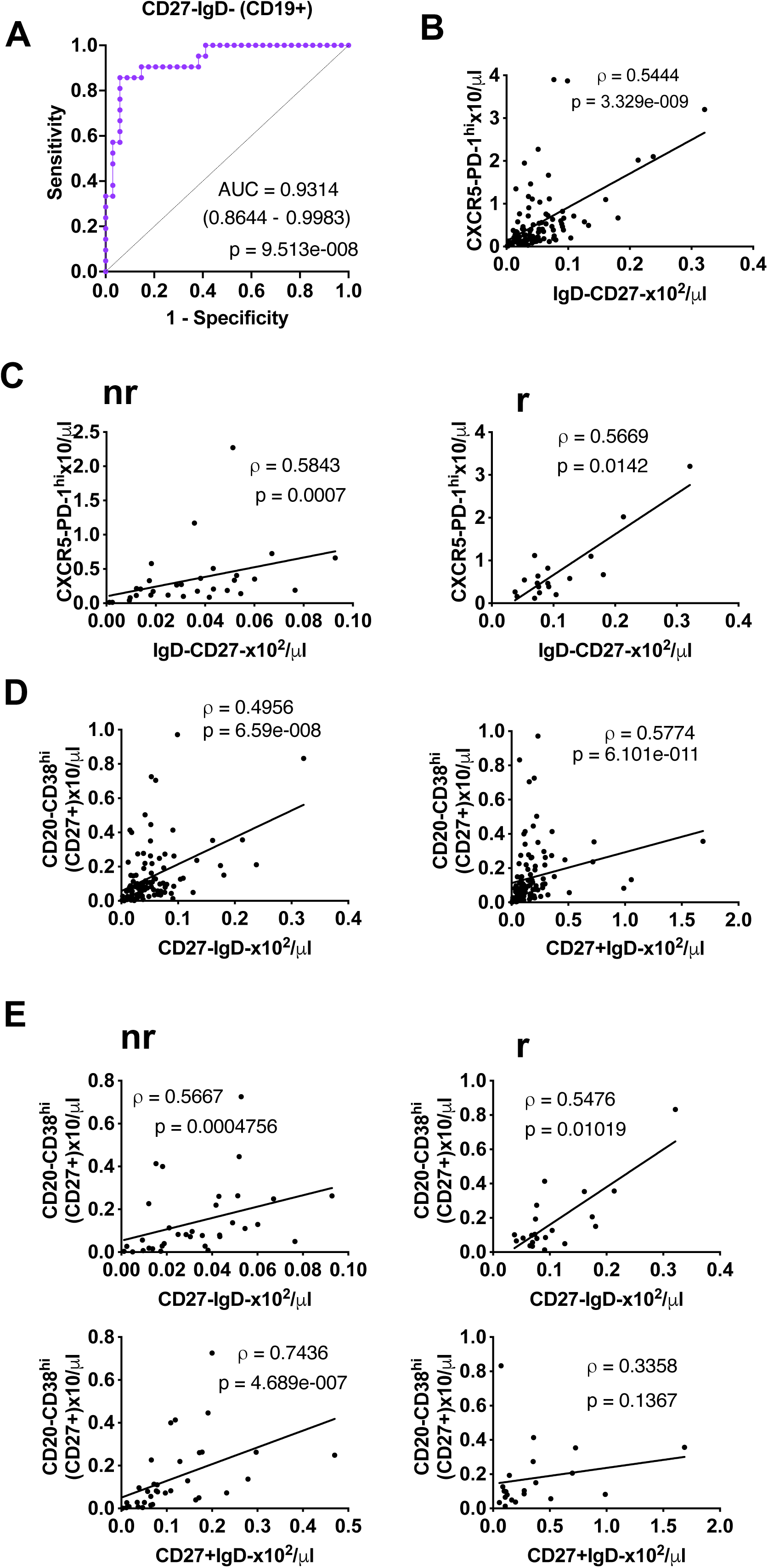
Baseline DN B-cell expansion predicts relapse and reveals distinct immune-cell association patterns according to clinical outcome. (A) Receiver operating characteristic (ROC) curve analyzing baseline absolute DN B-cell counts (CD19+CD27-IgD-) for predicting relapse within 12 months after diagnosis, in the cohort with available 1-year clinical outcome data (n=55; 34 non-relapsers and 21 relapsers). (B) Spearman correlation between baseline absolute DN B-cell and circulating peripheral helper T (cTph) cell counts in the total baseline nGCA cohort (n=102). (C) Correlation between baseline absolute DN B-cell and cTph cell counts stratified by 1-year clinical outcome into non-relapsers (n=34) and relapsers (n=21). (D) Correlation of baseline plasmablast counts (CD19+CD27+CD20-CD38^hi^) with absolute DN B-cell (left) and switched memory B-cell (CD19+CD27+IgD-) counts (right) in the total cohort (n=102). (E) Correlation of baseline plasmablast counts with absolute DN B-cell (upper) and switched memory B-cell counts (lower panels), stratified into non-relapsers (n=34, left) and relapsers (n=21, right). Statistical significance was assessed using Spearman’s rank correlation coefficient (ρ). **Alt text.** Baseline expansion of circulating double-negative B cells predicts relapse in newly diagnosed giant cell arteritis and is associated with distinct immune-cell interaction patterns. Double-negative B-cell counts correlate with circulating peripheral helper T cells and with plasmablast counts, while the relationship between plasmablasts and switched memory B cells is observed only in patients who remain relapse-free.

### Correlation Between DN B Cells and cTph Cells at Disease Onset

Given the established contribution of CD4+CXCR5-PD-1^hi^ peripheral helper T (cTph) cells to extrafollicular B-cell responses [24,25] and their emerging role in nGCA [4–6], we explored the possible association of circulating Tph (cTph) cells with circulating DN B cells at disease onset. Baseline absolute DN B cell counts showed a significant positive correlation with cTph cell counts (Fig. 5B). Notably, this association was maintained across the entire cohort and within both relapsing and non-relapsing patient subsets (Fig. 5C).

### Association Between Plasmablasts and B-Cell Precursors According to Relapse Status

We next investigated the association between baseline plasmablast counts and their putative precursor B-cell populations. Plasmablast counts correlated significantly with DN B-cell counts across the entire cohort (Fig. 5D), and this association persisted when relapsers and non-relapsers were analyzed separately (Fig. 5E). By contrast, although plasmablast counts also correlated with switched memory B cells in the overall cohort (Fig. 5D), this association was restricted to non-relapsers and not observed in relapsers (Fig. 5E).

## DISCUSSION

Giant cell arteritis (GCA) is a granulomatous large-vessel vasculitis of older adults, characterised by systemic inflammation and a risk of severe ischemic complications [1]. Despite therapeutic advances including interleukin-6-receptor blockade, treatment-related toxicity and disease relapse remain major clinical challenges [26]. A more precise understanding of the immunopathogenic mechanisms underlying disease persistence is therefore required to identify biomarkers capable of informing risk stratification and therapeutic decision-making [26].

Recent work in rheumatic diseases has emphasized that peripheral blood immune-cell profiling can identify clinically relevant cellular signatures associated with disease heterogeneity, treatment response, and relapse risk [27]. We herein show that patients with nGCA exhibit marked disturbances in circulating B-cell homeostasis, with reduced total CD19+ B-cell counts and an altered distribution of B-cell subsets. To our knowledge this is the first study to demonstrate increased absolute numbers of circulating DN B cells in nGCA and their association with relapse in patients receiving standard glucocorticoid monotherapy [15]. The inclusion of age- and sex-matched healthy controls strengthens the attribution of these immune alterations to disease-specific mechanisms. Moreover, the association between CD19+CD27-IgD-DN B cells and relapse risk underscores their biological relevance without further phenotypic subdivision [28].

The present work extends seminal observations of van der Geest and colleagues [10], who first demonstrated B-cell lymphopenia in active nGCA that tended to normalize during remission [10], suggesting that altered B-cell distribution is more closely related to inflammatory activity than to glucocorticoid exposure alone [10]. Consistent with this concept, our nGCA patients exhibited a significant rise in total B-cell counts over six months when analyzed as a single cohort. Additional studies have reported reduced baseline B-cell counts [11–14] and dynamic fluctuations in lymphocyte subsets during treatment [5,11], alongside persistent myeloid predominance [12], supporting altered immune cell trafficking and compartmentalization in this disease. Mechanistic investigations have further identified reduced frequencies of circulating CXCR3⁺ and CXCR5⁺ memory B-cells associated with increased levels of their ligands CXCL9 and CXCL13 both systemically and within vascular lesions [13], indicating active recruitment and organization of B-cells at sites of arterial inflammation [13].

Importantly, previous studies reporting reduced circulating B-cell numbers in nGCA did not stratify baseline counts according to subsequent clinical evolution under glucocorticoid therapy [10–14]. Consequently, the global reduction observed in earlier cohorts may have masked clinically relevant heterogeneity that becomes evident when patients are analyzed by 1-year outcome. In our cohort, decreased baseline B-cell numbers were confined to patients who remained relapse-free under standard therapy, whereas relapsers displayed counts exceeding HC.

Furthermore, expanding on findings that GCA B-cell lymphopenia reflects inflammation [10] and normalizes with improvement [10], we found that this dynamic pattern was restricted to non-relapsers. Relapsers maintaind elevated baseline B-cell counts and minimal longitudinal variation despite initial clinical improvement. This stability suggests a persistent, treatment-resistant pathogenic B-cell profile that predisposes to disease recurrence.

Indeed, baseline expansion of the DN subset emerged as the most distinctive feature of relapsers. Baseline DN B-cell counts independently predicted relapse risk after adjustment for inflammatory markers, suggesting that this subset reflects a pathogenic immune trajectory rather than a simple correlate of systemic inflammation. Our data align with evidence in systemic lupus erythematosus (SLE), where DN B-cell expansion correlates with nephritis, higher disease activity [29–31], and lower remission rates [30, 32]. Notably, Jenks et al. [33] identified an expanded DN B-cell subset as a distinct effector population and immediate precursor to antibody-secreting plasmablasts. While biopsy data were unavailable in our cohort due to an ultrasound-based diagnostic pathway, previous histopathological studies have demonstrated B-cell infiltration and tertiary lymphoid organ formation in GCA [7, 8]. This supports the possibility that the expansion of circulating DN B cells reported here reflects tissue-level extrafollicular immune responses.

Within this framework, expansion of the DN B-cell compartment in relapsing nGCA patients likely reflects a specialized immune phenotype poised to evade standard therapy. This evasion could be driven by a pathogenic synergy between a relative survival advantage of DN B cells under glucocorticoid pressure and preferential engagement of extrafollicular (EF) activation pathways. While glucocorticoids effectively target naïve and transitional B-cell populations, more differentiated or activated subsets, including DN B cells, may be comparatively less susceptible to these agents [34–36]. In parallel, recent evidence has challenged the traditional view of germinal centers as the exclusive drivers of autoimmunity, highlighting EF responses as a major source of pathogenic humoral activity [37,38]. In this context, DN B cells appear to be central components of the EF axis, integrating B-cell receptor (BCR), Toll-like receptor, cytokine and T-cell-derived signals [33,39]. These pathways promote the rapid differentiation of activated B-cells into plasmablasts and plasma cells [33], while operating outside the stringent tolerance checkpoints normally imposed within germinal centers [37, 38]. As a result, EF responses may favor the persistence and expansion of autoreactive clones [37, 38]. Maintenance of this alternative (“bypass”) activation circuit during glucocorticoid tapering may therefore contribute to incomplete resolution of vascular inflammation and increased susceptibility to relapse.

Additionally, expansion of DN B cells in nGCA patients who subsequently relapsed may reflect dysregulated interactions within a pathogenic extrafollicular T-cell-B-cell activation axis. A key observation from our study is the significant positive correlation between baseline DN B cells and circulating peripheral helper T (cTph) cells. While we previously reported an expansion of cTph cells in nGCA [6], their strong association here with the DN compartment provides a mechanistic framework linking two immune populations implicated in disease progression and relapse. As reviewed by He and Vinuesa [37], extrafollicular B-cell responses can be categorized into thymus-independent or thymus-dependent (TD) pathways, with the latter being particularly prominent in autoimmune conditions. Tph cells are recognized as potent drivers of these TD extrafollicular B-cell responses [24, 25], providing critical help through IL-21 and MAF signaling [25]. Notably, Bocharnikov et al. [25] demonstrated that Tph cells, which are markedly expanded in systemic autoimmunity, possess a unique capacity to promote memory B cell differentiation into plasmablasts, independently of CXCR5-mediated germinal center organization [25]. Furthermore, their work showed that Tph cells specifically correlate with the expansion of CD11c+ atypical B-cells, a population that overlaps significantly with the DN compartment [25]. A similar correlation of Tph cells with DN2 B cells was reported in childhood-onset SLE by Baxter et al. [31]. The universal correlation observed across our cohort suggests that the Tph-DN axis is a core component of the initial immune response in nGCA, likely facilitating rapid B-cell activation and effector differentiation at extrafollicular sites.

However, while this Tph-DN link was shared across our entire cohort, stratified analysis revealed that the downstream relationship between plasmablasts and upstream B-cell compartments diverged notably between non-relapsers and relapsers. In non-relapsers, baseline plasmablast counts correlated with both switched memory B cells and the DN subset, suggesting that plasmablast generation remained associated with both conventional memory responses and the DN compartment. Conversely, in relapsers, plasmablast levels correlated with DN B cells but showed no significant association with switched memory B cells. Although these findings do not establish direct lineage relationships, they raise the possibility that plasmablast generation in relapsers may be more closely linked to the DN B-cell compartment, whereas the contribution of the conventional switched-memory compartment is diminished. This interpretation is consistent with a greater contribution of extrafollicular-associated B-cell responses in patients who subsequently relapse. Such skewed extrafollicular pathways mirror the aggressive, treatment-resistant disease phenotypes observed when this axis is hyperactivated in SLE [25, 31].

Taken together, and paralleling emerging models of immune stratification in SLE [40, 41], our data suggest that nGCA may encompass distinct B-cell endotypes, with relapsers showing a more prominent DN/extrafollicular-associated profile and non-relapsers a more conventional germinal center-associated pattern.

Of note, B-cell counts were unaffected by ultrasound-defined disease patterns or concomitant PMR, which aligns with the current view of GCA as a continuous phenotypic spectrum rather than distinct cranial/extracranial entities [22, 26]. Indeed, approximately 35% of patients with traditional “cranial” symptoms exhibit large-vessel involvement upon imaging [42]. Moreover, the disease phenotype is dynamic; patients initially presenting with large-vessel GCA may develop cranial manifestations during follow-up, and vice versa [43,44].

Following current EULAR recommendations, patients were treated to balance clinical efficacy with glucocorticoid (GC) toxicity [15]. Because many patients achieve sustained remission with GC monotherapy and taper to ≤5 mg/day within one year [15, 45–47], upfront use of disease-modifying antirheumatic drugs (DMARDs) is reserved for selected cases to avoid unnecessary toxicity and costs [15, 48]. Our results suggest that baseline DN B-cell counts could refine this strategy by identifying patients who may benefit from early treatment intensification while sparing lower-risk individuals from overtreatment. A primary strength of this study lies in the integration of our fast-track US clinic with specialized laboratory infrastructure, ensuring the recruitment of treatment-naïve nGCA patients and the immediate processing of samples at diagnosis. This seamless coordination, coupled with rigorous clinical and laboratory follow-up, enhances the reliability of our longitudinal data. Furthermore, by stratifying patients based on one-year clinical outcomes, we identified immunological heterogeneities within the B-cell compartment that remain obscured in aggregate cohort analyses. Despite these strengths, several limitations warrant consideration. First, we did not perform functional assays; such experiments require large blood volumes, often difficult to obtain from acutely ill, elderly patients. Second, while our cohort size was sufficient to achieve statistical significance for our key observations, these findings represent a discovery phase for the relapse-predicting value of DN B-cell frequencies. Consequently, larger multicenter studies and external validation cohorts are required.

In summary, nGCA is characterized by marked disturbances in B-cell homeostasis, with expansion of the DN B-cell compartment emerging as the feature most closely associated with relapse. DN B-cell quantification may therefore provide a clinically relevant biomarker for risk stratification and further supports a role for extrafollicular immune activation in disease persistence.

## Funding

This work was supported by Ministerio de Economía y Competitividad-Instituto de Salud Carlos III (MINECO/ISCIII) grant number FIS PI25/00234 (http://www.isciii.es) and by Fondo Europeo de Desarrollo Regional (FEDER). The funders had no role in study design, data collection and analysis, decision to publish, or preparation of the manuscript.

## Competing interests

None related to this article

## Patient and Public Involvement statement

Patients or the public WERE NOT involved in the design, or conduct, or reporting, or dissemination plans of our research

## AUTHOR CONTRIBUTIONS

All authors were involved in drafting the article or revising it critically for important intellectual content, and all authors approved the final version to be published. Dr. Miranda-Carús had full access to all of the data in the study and takes responsibility for the integrity of the data and the accuracy of the data analysis.

**Study conception and design**. Miranda-Carús

**Acquisition of data**. Monjo-Henry, Nieto-Carvalhal, Campos-Fabre, García-Carazo, Villalba, Peiteado, de Miguel, Miranda-Carús

**Analysis and interpretation of data**. Monjo-Henry, Nieto-Carvalhal, Campos-Fabre, García-Carazo, de Miguel, Peiteado, Villalba, Miranda-Carús

**Wrote the manuscript**. Miranda-Carús

## DATA AVAILABILITY STATEMENT

The raw data supporting the conclusions of this article will be made available by the corresponding author upon reasonable request.

---

**Figure S1.**
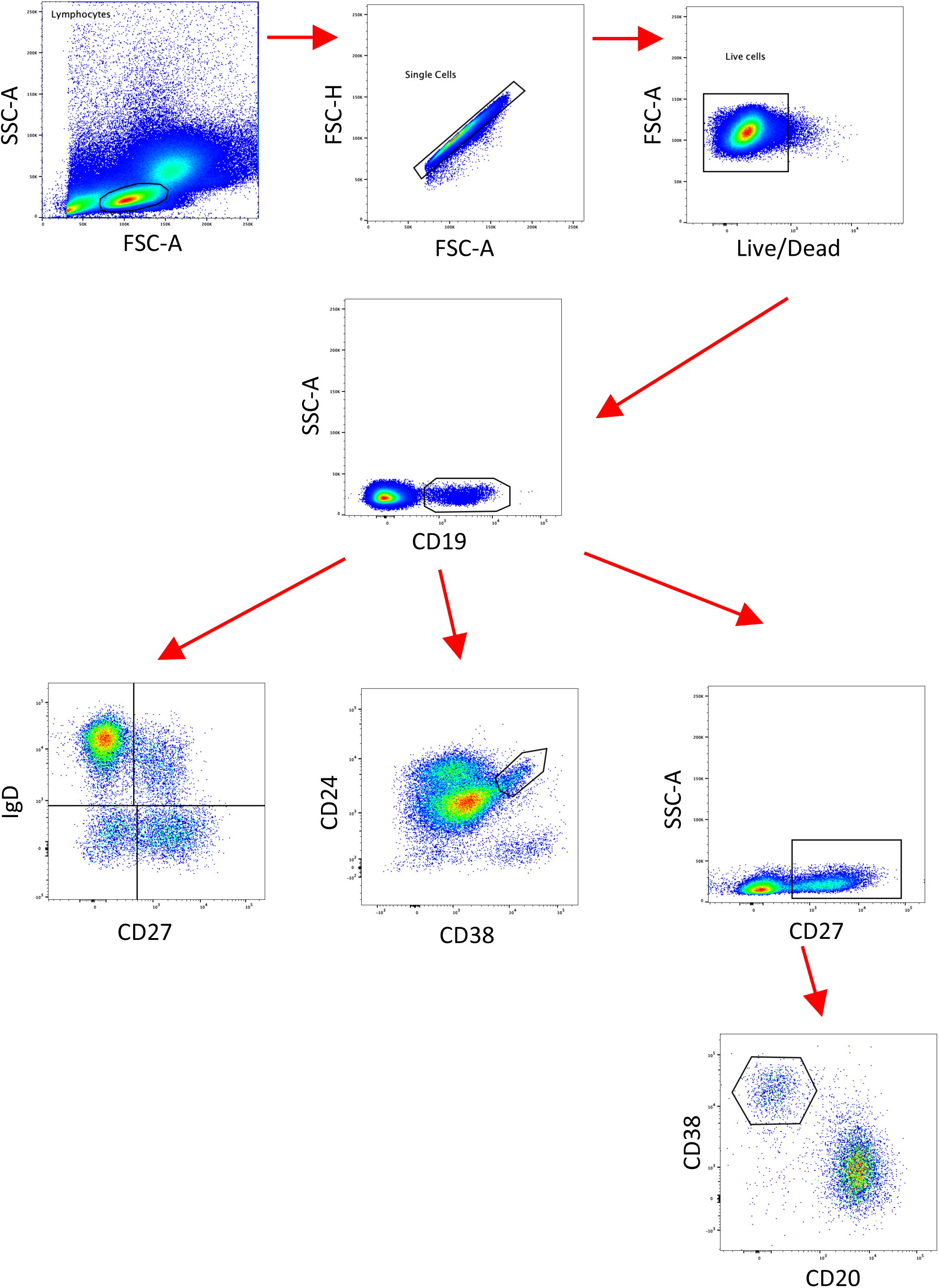
Gating strategy for the identification of circulating B cell subpopulations

**Figure S2.**
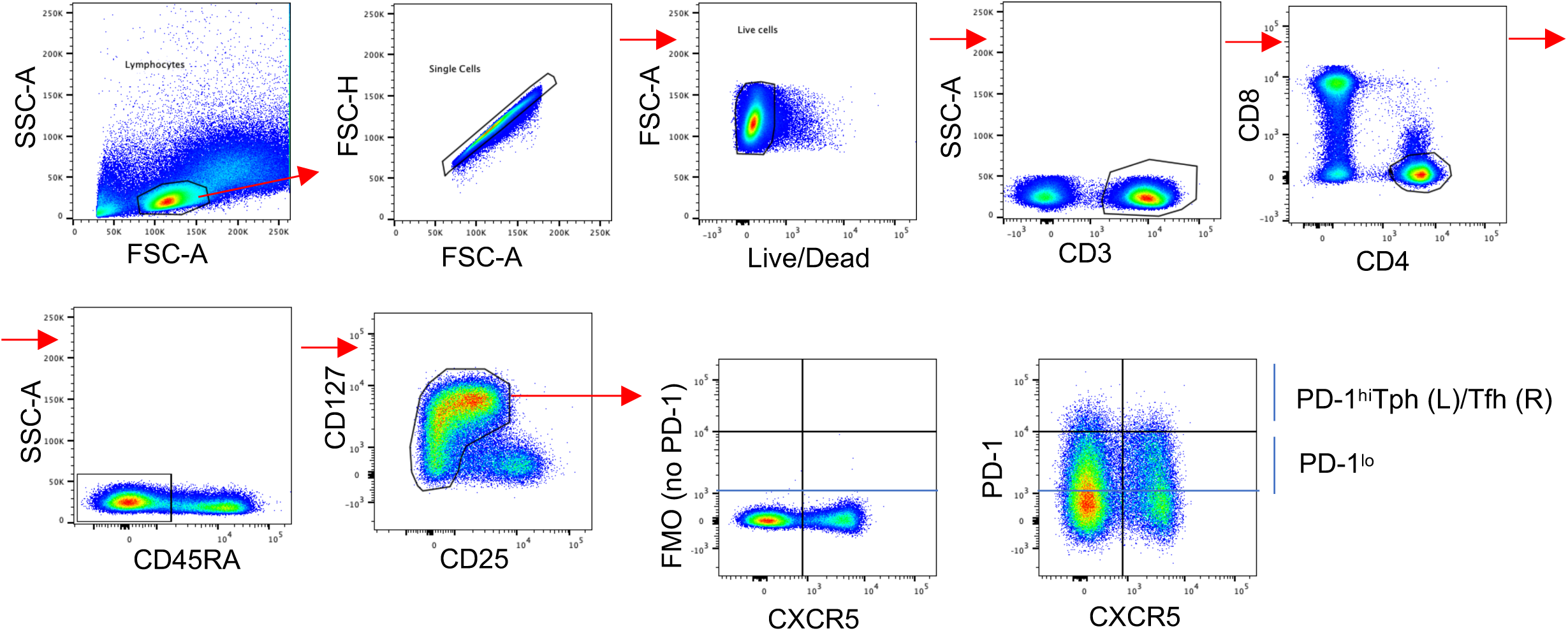
Gating strategy for the identification of circulating Tph cells. Adapted from Nuño L. et al, 2026 [Rheumatology (Oxford) 2026, Feb 4;65(2):keaf648.]

## Notes

### Competing Interest Statement

The authors have declared no competing interest.

## REFERENCES

1. Weyand CM, Goronzy JJ. Medium- and large-vessel vasculitis. N Engl J Med. 2003;349(2):160–9. doi: 10.1056/NEJMra022694.

2. Weyand CM, Goronzy JJ. Immunology of Giant Cell Arteritis. Circ Res. 2023;132(2):238–250. doi: 10.1161/CIRCRESAHA.122.322128.

3. Robinette ML, Rao DA, Monach PA. The Immunopathology of Giant Cell Arteritis Across Disease Spectra. Front Immunol. 2021;12:623716. doi: 10.3389/fimmu.2021.623716.

4. Desbois AC, Régnier P, Quiniou V, Lejoncour A, Maciejewski-Duval A, Comarmond C, et al. Specific Follicular Helper T Cell Signature in Takayasu Arteritis. Arthritis Rheumatol. 2021;73(7):1233–1243. doi: 10.1002/art.41672.

5. Matsumoto K, Suzuki K, Yoshimoto K, Seki N, Tsujimoto H, Chiba K, Takeuchi T. Significant association between clinical characteristics and changes in peripheral immuno-phenotype in large vessel vasculitis. Arthritis Res Ther. 2019;21(1):304. doi: 10.1186/s13075-019-2068-7.

6. Monjo-Henry I, Nieto-Carvalhal B, Uyaguari M, García-Carazo S, Balsa A, de Miguel E, Miranda-Carús ME. Circulating PD-1^hi^ CXCR5- and CXCR5+ CD4 T cells are elevated in patients with newly diagnosed giant cell arteritis, and predict relapse. Rheumatology (Oxford). 2025;64(6):3996–4004. doi: 10.1093/rheumatology/keaf001.

7. Ciccia F, Rizzo A, Maugeri R, Alessandro R, Croci S, Guggino G, et al. Ectopic expression of CXCL13, BAFF, APRIL and LT-β is associated with artery tertiary lymphoid organs in giant cell arteritis. Ann Rheum Dis 2017;76:235–43. doi: 10.1136/annrheumdis-2016-209217.

8. Graver JC, Boots AMH, Haacke EA, Diepstra A, Brouwer E, Sandovici M. Massive B cell infiltration and organization into artery tertiary lymphoid organs in the aorta of large vessel giant cell arteritis. Front Immunol 2019;10:83. doi: 10.3389/fimmu.2019.00083

9. Graver JC, Jiemy WF, Altulea DHA, van Sleen Y, Xu S, van der Geest KSM, et al. Cytokine producing B cells and their capability to polarize macrophages in giant cell arteritis. J Autoimmun. 2023;140:103111. doi: 10.1016/j.jaut.2023.103111.

10. van der Geest KSM, Abdulahad WH, Chalan P, Rutgers A, Horst G, Huitema MG, et al. Disturbed B cell homeostasis in newly diagnosed giant cell arteritis and polymyalgia rheumatica. Arthritis Rheumatol 2014;66:1927–38. doi: 10.1002/art.38625.

11. Kuret T, Frank-Bertoncelj M, Lakota K, Žigon P, Thallinger GG, Kopitar AN, Čučnik S, Tomšič M, Hočevar A, Sodin-Šemrl S. From Active to Non-active Giant Cell Arteritis: Longitudinal Monitoring of Patients on Glucocorticoid Therapy in Combination With Leflunomide. Front Med (Lausanne). 2022;8:827095. doi: 10.3389/fmed.2021.827095.

12. van Sleen Y, Graver JC, Abdulahad WH, van der Geest KSM, Boots AMH, Sandovici M, et al. Leukocyte dynamics reveal a persistent myeloid dominance in giant cell arteritis and polymyalgia rheumatica. Front Immunol. 2019;10:1981. doi: 10.3389/fimmu.2019.01981.

13. Graver JC, Abdulahad W, van der Geest KSM, Heeringa P, Boots AMH, Brouwer E, Sandovici M. Association of the CXCL9-CXCR3 and CXCL13-CXCR5 axes with B cell trafficking in giant cell arteritis and polymyalgia rheumatica. J Autoimmun. 2021;123:102684. doi: 10.1016/j.jaut.2021.102684.

14. Palamidas DA, Papadaki M, Paschalidis N, Pavlos E, Chatzis L, Palla P, et al. Multiparametric analysis in the peripheral blood of Giant Cell Arteritis and Polymyalgia Rheumatica patients at the early phases of steroid treatment reveals changes in cell subpopulations and lipid mediators: a preliminary study. Front Immunol. 2025;16:1594263. doi: 10.3389/fimmu.2025.1594263

15. Hellmich B, Agueda A, Monti S, Buttgereit F, de Boysson H, Brouwer E, et al. 2018 Update of the EULAR recommendations for the management of large vessel vasculitis. Ann Rheum Dis. 2020;79:19–30. doi: 10.1136/annrheumdis-2019-215672.

16. Chrysidis S, Duftner C, Dejaco C, Schäfer VS, Ramiro S, Carrara G, et al. Definitions and reliability assessment of elementary ultrasound lesions in giant cell arteritis: a study from the OMERACT large vessel vasculitis ultrasound Working group. RMD Open 2018;4:e000598. doi: 10.1136/rmdopen-2017-000598.

17. Schäfer VS, Juche A, Ramiro S, Krause A, Schmidt WA. Ultrasound cut-off values for intima-media thickness of temporal, facial and axillary arteries in giant cell arteritis. Rheumatology (Oxford) 2017;56:1479–83. doi: 10.1093/rheumatology/kex143.

18. De Miguel E, Beltran LM, Monjo I, et al. Atherosclerosis as a potential pitfall in the diagnosis of giant cell arteritis. Rheumatology (Oxford) 2018;57:318–21. doi: 10.1093/rheumatology/kex381.

19. Ješe R, Rotar Ž, Tomšič M, Hočevar A. The cut-off values for the intima-media complex thickness assessed by colour Doppler sonography in seven cranial and aortic arch arteries. Rheumatology (Oxford). 2021 Mar 2;60(3):1346–1352. doi: 10.1093/rheumatology/keaa578.

20. Monjo-Henry I, Fernández-Fernández E, Mostaza JM, Lahoz C, Molina-Collada J, de Miguel E. Ultrasound halo count in the differential diagnosis of atherosclerosis and large vessel giant cell arteritis. Arthritis Res Ther 2023;25:23. doi: 10.1186/s13075-023-03002-0.

21. De Miguel E, Macchioni P, Conticini E, Campochiaro C, Karalilova R, Monti S, et al. Rheumatology (Oxford). 2024;63:158–164. doi: 10.1093/rheumatology/kead189. PMID: 37129541.

22. Dejaco C, Ramiro S, Bond M, Bosch P, Ponte C, Mackie SL, et al. EULAR recommendations for the use of imaging in large vessel vasculitis in clinical practice: 2023 update. Ann Rheum Dis. Ann Rheum Dis. 2024 May 15;83(6):741–751. doi: 10.1136/ard-2023-224543.

23. Nuño L, Nieto-Carvalhal B, Gutiérrez-Riart M, Novella-Navarro M, Peiteado D, Monjo I, et al. Expanded blood PD-1hi Tfh cells in seropositive clinically suspect arthralgia predict progression to rheumatoid arthritis. Rheumatology (Oxford). 2026;65(2):keaf648. doi: 10.1093/rheumatology/keaf648. PMID: 41346250.

24. Rao DA, Gurish MF, Marshall JL, Slowikowski K, Fonseka CY, Liu Y, et al. Pathologically expanded peripheral T helper cell subset drives B cells in rheumatoid arthritis. Nature. 2017;542(7639):110–114. doi: 10.1038/nature20810.

25. Bocharnikov AV, Keegan J, Wacleche VS, Cao Y, Fonseka CY, Wang G, et al. Accelerating Medicines Partnership (AMP) RA/SLE Network. PD-1hiCXCR5-T peripheral helper cells promote B cell responses in lupus via MAF and IL-21. JCI Insight. 2019;4(20):e130062. doi: 10.1172/jci.insight.130062.

26. Tomelleri A, van der Geest KSM, Sebastian A, van Sleen Y, Schmidt W, Dejaco C, et al. Disease stratification in giant cell arteritis to reduce relapses and prevent long-term vascular damage. The Lancet Rheumatology 2021;3:e886–95. doi: 10.1016/S2665-9913(21)00277-0.

27. Rao DA. Immune-cell profiling to guide stratification and treatment of patients with rheumatic diseases. Nat Rev Rheumatol. 2025;21(11):657–670. doi: 10.1038/s41584-025-01291-0.

28. Sanz I, Wei C, Lee FE, Anolik J. Phenotypic and functional heterogeneity of human memory B cells. Semin Immunol. 2008; 20:67–82. doi: 10.1016/j.smim.2007.12.006.

29. Wei C, Anolik J, Cappione A, Zheng B, Pugh-Bernard A, Brooks J, et al. A new population of cells lacking expression of CD27 represents a notable component of the B cell memory compartment in systemic lupus erythematosus. J Immunol. 2007;178(10):6624–33. doi: 10.4049/jimmunol.178.10.6624.

30. You X, Zhang R, Shao M, He J, Chen J, Liu J, et al. Double Negative B Cell Is Associated With Renal Impairment in Systemic Lupus Erythematosus and Acts as a Marker for Nephritis Remission. Front Med (Lausanne). 2020;7:85. doi: 10.3389/fmed.2020.00085.

31. Baxter RM, Wang CS, Garcia-Perez JE, Kong DS, Coleman BM, Larchenko V, et al. Expansion of extrafollicular B and T cell subsets in childhood-onset systemic lupus erythematosus. Front Immunol. 2023;14:1208282. doi: 10.3389/fimmu.2023.1208282.

32. Wangriatisak K, de Vries C, Sharma RK, Huang W, Grönwall C, Pisitkun P, et al. Association between peripheral activated naive and double negative 2 B-cell subsets and clinical parameters in lupus nephritis patients. Scand J Immunol. 2025;101(1):e13427. doi: 10.1111/sji.13427.

33. Jenks SA, Cashman KS, Zumaquero E, Marigorta UM, Patel AV, Wang X, et al. Distinct Effector B Cells Induced by Unregulated Toll-like Receptor 7 Contribute to Pathogenic Responses in Systemic Lupus Erythematosus. Immunity. 2018 Oct 16;49(4):725–739.e6. doi: 10.1016/j.immuni.2018.08.015

34. Cain DW, Cidlowski JA. Immune regulation by glucocorticoids. Nat Rev Immunol. 2017;17(4):233–247. doi: 10.1038/nri.2017.1.

35. Franco LM, Gadkari M, Howe KN, Sun J, Kardava L, et al. Immune regulation by glucocorticoids can be linked to cell type-dependent transcriptional responses. J Exp Med. 2019;216(2):384–406. doi: 10.1084/jem.20180595.

36. Almohammad K, Young M, Vettorazzi S, Greulich F, Alkhatib M, Tuckermann J, Jumaa H, Setz CS. IgD-Expressing Mature B Cells Exhibit Enhanced Sensitivity to Glucocorticoid-Induced Cell Death. Eur J Immunol. 2026;56:e70137. doi: 10.1002/eji.70137.

37. He Y, Vinuesa CG. Germinal center versus extrafollicular responses in systemic autoimmunity: Who turns the blade on self? Adv Immunol. 2024;162:109–133. doi: 10.1016/bs.ai.2024.02.002.

38. Eisenbarth SC, Batista F, Cyster J, Elsner R, Kelsoe G, Lund FE, et al. A roadmap for defining “extrafollicular” B cell responses. Immunity. 2025;58(11):2627–2645. doi: 10.1016/j.immuni.2025.08.007.

39. Staniek J, Rizzi M. Signaling Activation and Modulation in Extrafollicular B Cell Responses. Immunol Rev. 2025;330(1):e70004. doi: 10.1111/imr.70004.

40. Suurmond J, Atisha-Fregoso Y, Barlev AN, Calderon SA, Mackay MC, Aranow C, Diamond B. Patterns of ANA+ B cells for SLE patient stratification. JCI Insight. 2019;4(9):e127885. doi: 10.1172/jci.insight.127885.

41. Faliti CE, Van TTP, Anam FA, Cheedarla N, Williams ME, Mishra AK, et al. Disease-associated B cells and immune endotypes shape adaptive immune responses to SARS-CoV-2 mRNA vaccination in human SLE. Nat Immunol. 2025;26(1):131–145. doi: 10.1038/s41590-024-02010-9.

42. Prieto-González S, Depetris M, García-Martínez A, Espígol-Frigolé G, Tavera-Bahillo I, Corbera-Bellata M, et al. Positron emission tomography assessment of large vessel inflammation in patients with newly diagnosed, biopsy-proven giant cell arteritis: a prospective, case-control study. Ann Rheum Dis 2014;73:1388–92. doi:10.1136/annrheumdis-2013-204572.

43. Narváez J, Estrada P, López-Vives L, Ricse M, Zacarías A, Heredia S, et al. Prevalence of ischemic complications in patients with giant cell arteritis presenting with apparently isolated Polymyalgia Rheumatica. Semin Arthritis Rheum 2015;45:328–33. doi: 10.1016/j.semarthrit.2015.06.009.

44. Fernández-Fernández E, Monjo I, Peiteado D, Balsa A, De Miguel E. Validity of the EULAR recommendations on the use of ultrasound in the diagnosis of giant cell arteritis. RMD Open 2022;8: e002120. doi: 10.1136/rmdopen-2021-002120.

45. Alba MA, García-Martínez A, Prieto-González S, Tavera-Bahillo I, Corbera-Bellalta M, Planas-Rigol E, et al. Relapses in patients with giant cell arteritis: prevalence, characteristics, and associated clinical findings in a longitudinally followed cohort of 106 patients. Medicine (Baltimore) 2014;93:194–201. doi: 10.1097/MD.0000000000000033.

46. Kermani TA, Warrington KJ, Cuthbertson D, Carette S, Hoffman GS, Khalidi NA, et al. Disease relapses among patients with giant cell arteritis: a prospective, longitudinal cohort study. J Rheumatol 2015;42:1213–7. doi: 10.3899/jrheum.141347.

47. Martinez-Lado L, Calviño-Díaz C, Piñeiro A, Dierssen T, Vazquez-Rodriguez TR, Miranda-Filloy JA, et al. Relapses and recurrences in giant cell arteritis: a population-based study of patients with biopsy-proven disease from northwestern Spain. Medicine (Baltimore) 2011;90:186–93. doi: 10.1097/MD.0b013e31821c4fad.

48. Jogimahanti AV, Kini AT, Irwin LE, Lee AG. The cost-effectiveness of tocilizumab (Actemra) therapy in giant cell arteritis. J. Neuroophthalmol. 2021;41:342–350 doi: 10.1097/WNO.0000000000001220.

